# Efficiently inferring the demographic history of many populations with allele count data

**DOI:** 10.1101/287268

**Authors:** John A. Kamm, Jonathan Terhorst, Richard Durbin, Yun S. Song

## Abstract

The sample frequency spectrum (SFS), or histogram of allele counts, is an important summary statistic in evolutionary biology, and is often used to infer the history of population size changes, migrations, and other demographic events affecting a set of populations. The expected multipopulation SFS under a given demographic model can be efficiently computed when the populations in the model are related by a tree, scaling to hundreds of populations. Admixture, back-migration, and introgression are common natural processes that violate the assumption of a tree-like population history, however, and until now the expected SFS could be computed for only a handful of populations when the demographic history is not a tree. In this article, we present a new method for efficiently computing the expected SFS and linear functionals of it, for demographies described by general directed acyclic graphs. This method can scale to more populations than previously possible for complex demographic histories including admixture. We apply our method to an 8-population SFS to estimate the timing and strength of a proposed “basal Eurasian” admixture event in human history. We implement and release our method in a new open-source software package `momi2`.

## 1 Introduction

All natural populations undergo evolutionary processes of migration, size changes, and divergence, and the history of these demographic events shape their present genetic diversity. Thus, inferring demographic history is of central concern in evolutionary and population genetics, both for its intrinsic interest (e.g., in dating the out-of-Africa migration of modern humans (Schaffner et al., 2005; Gutenkunst et al., 2009)) and also for biological applications (such as distinguishing the effects of natural selection from demography (Beaumont and Nichols, 1996; Boyko et al., 2008)). However, genetic sequence data and the space of possible demographic models are both very high dimensional objects, leading to numerous statistical and computational challenges when inferring demographic history from genetic data.

The joint sample frequency spectrum (SFS) is the multidimensional histogram of mutant allele counts in a sample of DNA sequences, and is a popular summary statistic which lies at the core of thousands of empirical studies in population genetics; e.g., see Wakeley and Hey (1997); Griffiths and Tavaré (1998); Nielsen (2000); Gutenkunst et al. (2009); Coventry et al. (2010); Gazave et al. (2014); Gravel et al. (2011); Nelson et al. (2012); Excoffier et al. (2013); Jenkins et al. (2014); Bhaskar et al. (2015); Jouganous et al. (2017). Recently, progress has also been made on theoretical fronts to characterize statistical properties of SFS-based inference. In particular, studies of identifiability (Myers et al., 2008; Bhaskar and Song, 2014) and the rate of convergence (Terhorst and Song, 2015; Baharian and Gravel, 2018) have been carried out.

Demographic history can be inferred by fitting the observed value of the SFS to its expected value in a composite likelihood framework. The expected SFS can be efficiently computed when the demographic history is a tree, and in previous work we developed a method *momi* to compute the SFS of hundreds of populations related by a tree (Kamm et al., 2017). However, natural populations are often related by a more complex history that is not tree-like, as gene flow (the exchange of migrants between populations) adds extra edges to the topology associated with the demographic history. In this case, computing the expected SFS is much more computationally demanding, and existing methods for computing the exact expected SFS can scale to only a handful of populations (Gutenkunst et al., 2009; Jouganous et al., 2017).

In this article, we extend our previous algorithm *momi* to handle discrete (or pulse) migration events between populations, in which case demographies are described by general directed acyclic graphs (DAGs). Our new method `momi2` can compute the *exact* expected SFS with admixture for more populations than previously possible, and uses novel insights from a stochastic process known as the Lookdown Construction (Donnelly and Kurtz, 1996; Donnelly et al., 1999). In addition, `momi2` utilizes automatic differentiation (Corliss et al., 2002; Bhaskar et al., 2015) to compute gradients of the SFS, which we use to efficiently search the parameter space during optimization. Finally, `momi2` can efficiently compute linear functionals of the SFS, which we exploit to compute the expected values of a number of standard population genetic summary statistics under complex demography.

The rest of this paper as organized as follows. In Section 2 we provide some background and survey related work. Section 3 describes our method, first with an illustrative example in Section 3.1, and then with formulas and pseudocode in Section 3.2. Finally, in Section 4, we apply our method to an 8-population SFS, including ancient and contemporary human populations, to estimate the timing and strength of a proposed “basal Eurasian” admixture event in human history. We defer all proofs to the Appendix.

## 2 Background

Suppose a sample of **n** = (*n*_1_, …, *n*_*𝒟*_) genomes have been sampled from 𝒟 “demes” or populations. The positions in the genome where the samples are not all identical are called *segregating sites*. In most organisms mutations are rare; most sites are not segregating. It is therefore reasonable to assume, as we do from now on, that each position in the genome has experienced at most a single mutation in its history, and that each individual can be labeled as having the “ancestral” or “derived” (mutant) allele at each segregating site. In population genetics, this simplifying assumption is known as the *infinite sites model*.

The *sample frequency spectrum* (SFS) is a -dimensional array 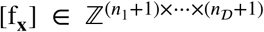 whose entry f_**x**_ counts the number of segregating sites with exactly **x** copies of the derived allele and **n** − **x** copies of the ancestral allele, where 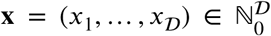 with 0 ≤ *x*_*d*_ ≤ *n*_*d*_ for each *d* = 1, …, 𝒟. Note we only consider segregating sites with 2 alleles, so f_**0**_ = f_**n**_ = 0 by definition. Compared to the full data set (i.e., the complete genetic sequences of all *n* = *n*_1_ + … + *n*_*d*_ genomes), the SFS [f_**x**_] is a compressed, low-dimensional summary which nevertheless preserves much of the signal about the various population size changes, divergence times, and admixture events that occurred over the course of the populations’ history.

### 2.1 Demographic events

The expected multipopulation SFS can be obtained by integrating over random genealogies formed by a backwards-in-time stochastic process known as the *structured coalescent* (Kingman, 1982; Takahata, 1988; Notohara, 1990). Before getting to the technical details, we first review the basic dynamics of this process in order to build intuition for how the data have power to infer population splits, size changes, and gene flow events. See Durrett (2008) for a more detailed introduction.

Informally, the topology and branch lengths of genealogies are affected by a demographic history in two ways:

1. Two lineages may not coalesce into a common ancestor until they reside in the same population, and the time until this occurs is affected by migration patterns and population split times.
2. At any particular point in time, two members within the same population are more likely to have a common parent if the population size is small; so, for example, residents of a small village will typically be more closely related than residents of a large city.

Regarding the second point, we define the scaled *effective population size η* (t) such that the rate at which any two lineages find a common ancestor at time *t* is 1/*η*(*t*). Under the simplest random mating model (the Wright Fisher model, cf. Durrett (2008)), the census population size exactly equals *Tη*, where *T* is the number of generations per unit time; more generally, *η* scales with the number of breeding individuals in the population. Thus, estimating *η* allows us to infer size change events such as bottlenecks, exponential growth, and population crashes.

If we could observe the true distribution of genealogies, then we could directly infer demographic history from the waiting times between coalescence events, following the principles listed in items 1 and 2 above. However, since genealogies are never directly observed, we must make inferences about demographic history indirectly using mutation data.

### 2.2 Likelihoods and the site frequency spectrum

Consider a genome with *L* positions and mutation rate 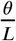 per position. So at any given position in the genome, mutations arise on the tree there as a Poisson point process with rate 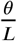. The chance of 2 or more mutations at a single position is 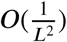, and taking the limit *L* → ∞ we arrive at the aforementioned *infinite sites approximation* (Kimura, 1969; Durrett, 2008), which assumes that each segregatin site was caused by asingle mutation, so that each allele may be labeled as ancestral or derived.

The observed segregating sites are not independent, because trees at neighboring positions are correlated. Unfortunately, even in the simplest case of a single population with constant size, an analytic expression for the likelihood of mutation data at a set of linked (non-independent) sites is not known (Bhaskar et al., 2015). Therefore, SFS data are generally used with composite likelihood methods. Recall that f_**x**_ is the total number of segregating sites with derived allele count pattern **x** = (*x*_1_,…, *x*_*𝒟*_). Define 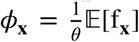, i.e., *Φ*_**x**_ is the expected frequency of **x** per unit mutation rate. Equivalently *Φ*_**x**_ is the expected branch length subtending **x** leaves in a random coalescent tree (i.e., with *x*_1_ descendants in population 1, *x*_2_ in population 2, and so on). A commonly used composite likelihood is the *Poisson random field* model (Sawyer and Hartl, 1992), which assumes that the total number of segregating sites is Poisson with rate *θ* Σ_**x**_*Φ*_**x**_;, and that the patterns at the observed sites are independent with sampling probabilities proportional to *Φ*_**x**_; this yields a log-likelihood of

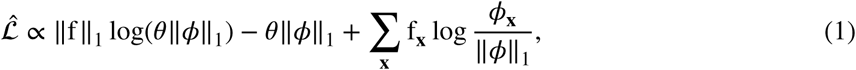

where ‖f‖_1_ = Σ_**x**_ f_**x**_ and ‖ *Φ*‖_1_ = Σ_**x**_ *Φ*_**x**_ Demographic history can then be inferred by searching for the parameter values that maximize 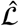.

### 2.3 Existing work and our contribution

The composite log-likelihood 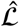 in (1) requires us to compute *Φ*_**x**_, the expected branch length subtending **x**. Let *G* be a random genealogical tree sampled under the demography, and *L*_**x**_(*G*) be the total length of all branches in *G* subtending **x**, so that

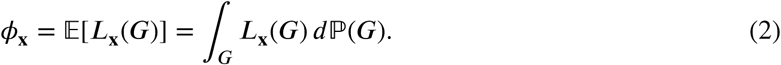

The integral (2) is difficult since the support of *G* is at least as large as the number of labeled binary trees with *n* leaves, a quantity which grows faster than exponentially in the sample size *n*. Consequently, a number of different methods have been proposed for evaluating (or approximating) the integral (2). Sampling-based methods include Markov chain Monte Carlo (Griffiths and Tavaré, 1997; Nielsen, 2000), importance sampling (Stephens and Donnelly, 2000; De Iorio and Griffiths, 2004), simulation (Excoffier and Foll, 2011; Excoffier et al., 2013), and ABC (Wegmann et al., 2010). The main advantage of these methods is their flexibility: since, as mentioned above, computing the likelihood of the data is trivial conditional on *G*, these methods can be used on a rich class of models. However, the high dimension of *Φ*_**x**_ makes it impractical to compute by sampling unless 𝒟 is small. In particular, since the support of **x** grows like *O*(*n*^*𝒟*^), Monte Carlo methods will assign zero mass to configurations that are actually observed in the data.

A second approach, implemented in the software *∂a∂i* (Gutenkunst et al., 2009), computes *Φ*_**x**_ by numerically solving PDEs arising from the Wright-Fisher diffusion (Ewens, 2004), which is dual to the coalescent process described above. For 𝒟 populations, this involves numerically solving a 𝒟-dimensional integral. The initial *∂a∂i* method in Gutenkunst et al. (2009) could handle up to 𝒟 = 3 populations; subsequent improvements (Lukić and Hey, 2012; Jouganous et al., 2017) extended this to 𝒟 = 4 and then 𝒟 = 5 populations by using spectral representations or alternative basis functions for solving the PDEs.

The third approach for computing *Φ*_**x**_, which includes our method, integrates over the sample allele frequencies “backwards-in-time”, exploiting conditional independence relationships to reduce computation. This involves considering the alleles of the sample’s ancestors at different points in the demographic history, and integrating out these random variables via inference algorithms for probabilistic graphical models (Pearl, 1982; Felsenstein, 1981; Lauritzen and Spiegelhalter, 1988; Koller and Friedman, 2009). Bryant et al. (2012) and Chen (2012) computed the SFS for finite- and infinite-sites models using this backward-in-time approach under the coalescent. De Maio et al. (2015) and Kamm et al. (2017) substantially lowered the computational burden of this approach by replacing the coalescent with the *continuous-time Moran model* (Durrett, 2008), a stochastic process which induces the same sampling distribution as the coalescent, but using a much smaller state space. However, until now these Moran-based approaches have been limited to analyzing tree-shaped demographies without admixture between populations.

The main contribution of this paper is to extend our previous Moran-based method (Kamm et al., 2017) to allow for demographies defined on general directed acyclic graphs (DAGs), thus allowing for admixture between populations. We describe and implement an algorithm for computing the expected infinite-sites SFS under the multipopulation Moran model with size changes, exponential growth, population splits, and point admixture events (i.e., instantaneous migration “pulses”). This substantially enlarges the space of demographies for which the expected SFS can be accurately computed.

Additionally, our algorithm computes not only individual SFS entries, but also linear functionals of the expected SFS. Specifically, our method computes rank-1 tensor products of the SFS in the same time as a single entry (general linear functionals are sums of these rank-1 products). A number of widely-used statistics in population genetics can be expressed as SFS functionals, and to our knowledge our method is the first to compute expectations of these statistics under complex demography.

We demonstrate our method by using it to infer the history of eight human subpopulations that have undergone multiple admixture events. We complement our theoretical contributions with an open-source, user-friendly software implementation that will enable practitioners to deploy our method. The software uses automatic differentiation (Corliss et al., 2002; Bhaskar et al., 2015; Maclaurin et al., 2015) to compute derivatives of the SFS, leading to efficient optimization and parameter inference. Our package, called momi2, is available for download at https://github.com/popgenmethods/momi2.

## 3 Method

In this section, we describe the algorithm implemented in `momi2` for computing the expected SFS under complex demographies. We begin in Subsection 3.1 with an illustrative example that highlights the novel aspects of our work. Then in Subsection 3.2 we provide pseudocode for our algorithm, and state the formulas used by our algorithm as Propositions. The proofs of these Propositions, and the proof for the correctness of our algorithm, requires substantial additional notation, and we defer this to Appendix A.2. For ease of reference, the symbols we use are summarized in Table 1.

**Table 1:**
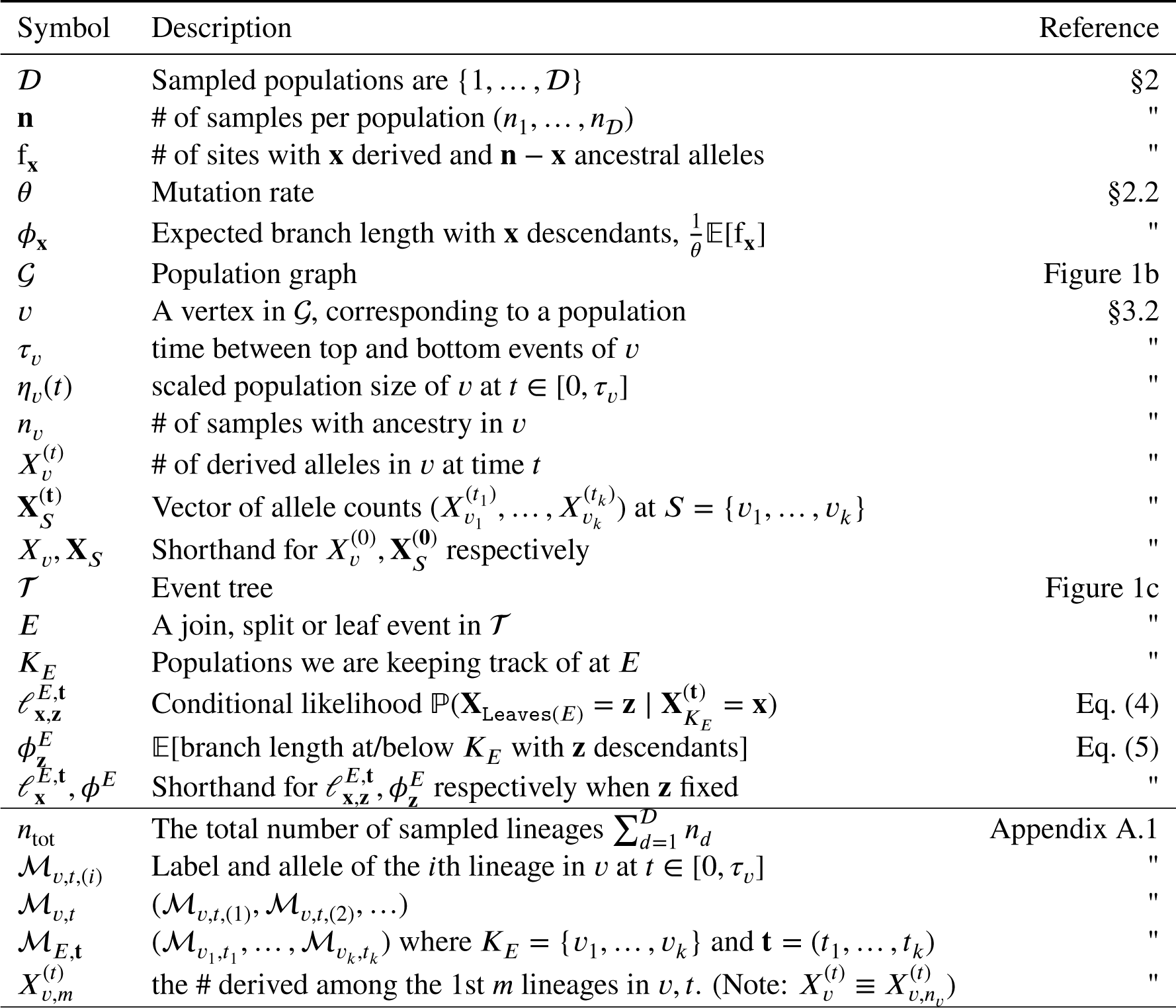
Notation. Rows in the second part of the table are only used for proofs in the Appendix and may be ignored in the main text.

### 3.1 Example

Consider the model depicted in Figure 1. This model has 3 sampled (leaf) populations, related as follows. Population 1 and 2 are sisters, with Population 3 an outgroup to them; however a pulse of migrants from 3 to 2 occurs after the spilt between populations 1 and 2.

**Figure 1:**
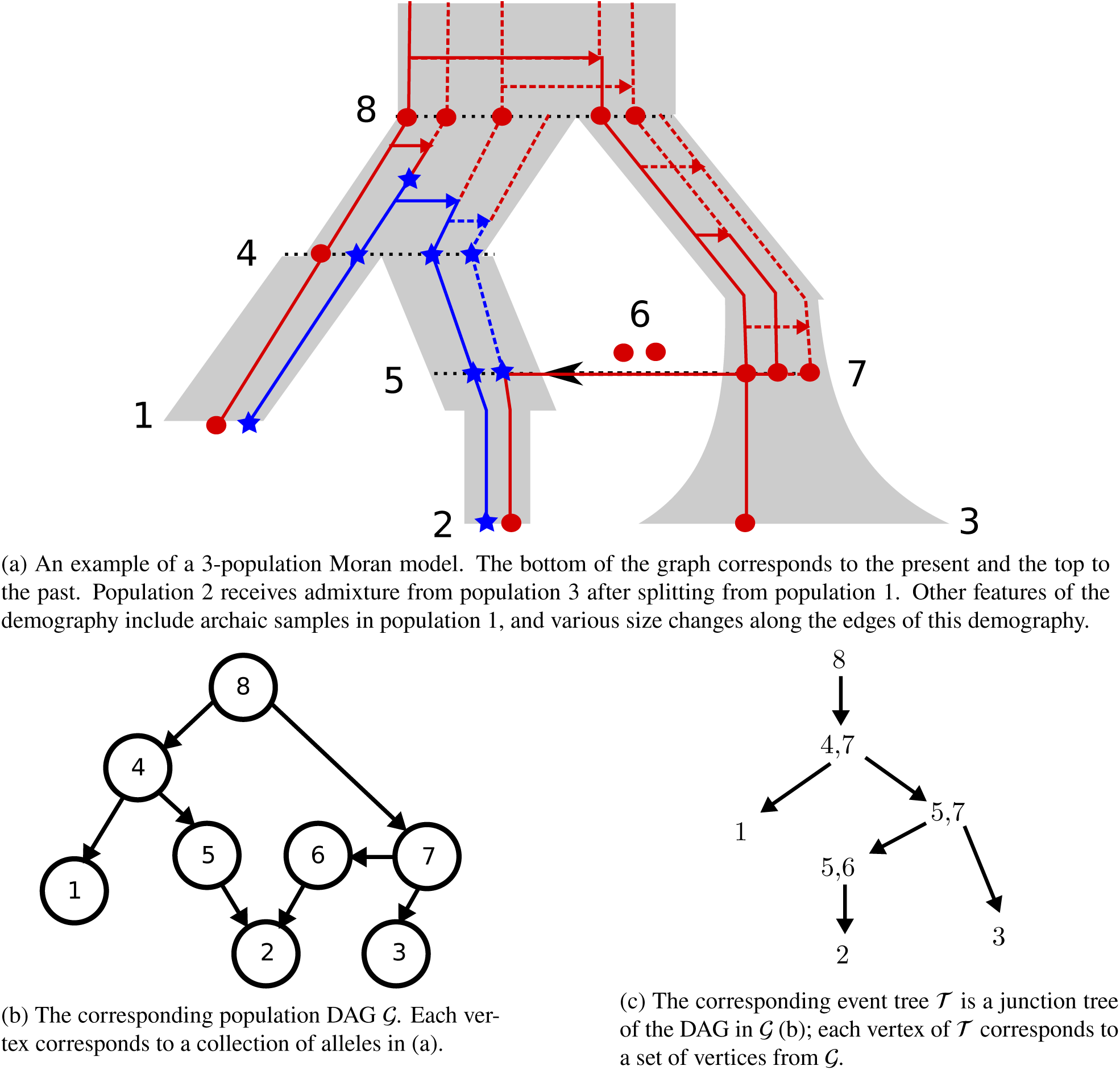
An illustrative example of a 3-population Moran model (a), along with (b) the corresponding DAG 𝒢, and (c) event tree 𝒯.

At the leaves, we obersve (*n*_1_, *n*_2_, *n*_3_) = (2, 2, 1) samples, with (*X*_1_, *X*_2_, *X*_3_) = (1,1,0) copies of the derived (blue star) allele. We wish to compute the expected number of mutations with pattern (*X*_1_, *X*_2_, *X*_3_); to do this, we will integrate over the unobserved variables (*X*_4_, *X*_5_, *X*_6,_ *X*_7_, *X*_8_) which which represent allele counts at certain internal positions within the demography. The random variables *X*_1,_ *X*_8_ are related to each other by the DAG 𝒢 in Figure 1b, with an edge from population *v* to *w* if alleles may pass directly from *v* into *w*.

In the realization of Figure 1, the hidden blue allele counts are (*X*_4_, *X*_5_, *X*_6_, *X*_7_, *X*_8_) = (3, 2, 0, 0, 0). The blue mutation occurs in the edge above *X*_4_, spreading to 3 of the lineages. One copy of the blue allele moves on to *X*_1_, while 2 copies move on to *X*_5_. However, due to the admixture, *X*_2_ only inherits 1 blue allele from *X*_5_, inheriting a red allele from the 2 red alleles at *X*_6_.

Under the infinite-sites assumption described above, the mutation observed at this site arose at a single point in the genealogical tree depicted in Figure 1. To compute the expected number of mutations with (*X*_1_, *X*_2_, *X*_3_) = (1, 1, 0), we may condition on the population (i.e., the edge in Figure 1) on which it arose:

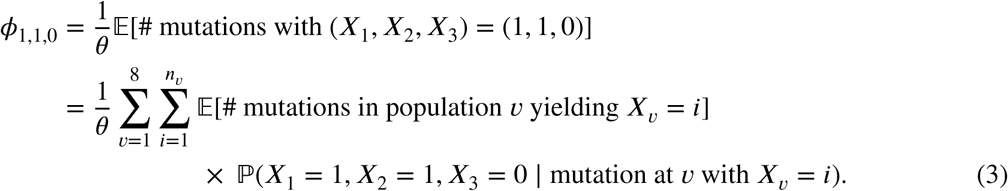

We call 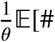 mutations at *v* with *X*_*v*_ = *i*], the “truncated SFS”; this only concerns events within a single population and can be computed using the method `momi1`. We refer the interested reader to (Kamm et al., 2017) for further details.

The second term ℙ(*X*_1_ = 1, *X*_2_ = 1, *X*_3_ = 0 | mutation at *v* with *X*_*v*_ = *i*) gives the conditional likelihood of observing the data given a mutation and its allele count at population *v*. In `momi1`, we computed this term in the case where 𝒢 is a tree without admixture using the sum-product (also known as belief propagation) algorithm (Felsenstein, 1981; Koller and Friedman, 2009).

The main result of this work is to extend our previous dynamic program to the case where 𝒢 is given by a DAG due to admixture. We will use a dynamic program that is essentially a kind of junction tree algorithm (Koller and Friedman, 2009). This algorithm works by decomposing a DAG graph into a tree in such a way that vertices in the tree correspond to collections of nodes in the original graph. Belief propagation is then applied to the tree decomposition.

We illustrate our algorithm using the example demography in Figure 1c. We call the tree decomposition 𝒯 an *event tree*, because each internal node corresponds to either an admixture or split event.^1^ We construct the event tree 𝒯, and compute the conditional likelihoods ℙ(*X*_1_ = 1, *X*_2_ = 1, *X*_3_ = 0 | mutation at *v* with *X*_*v*_ = *i*),as follows:

1. We initially start with a collection of 3 singleton sets of leaf populations: {{1}, {2}, {3}}. For *v* = 1, 2, 3, we also keep track of conditional likelihoods of the data beneath *v*; since we are at the leaves, these are simply the Kronecker delta functions

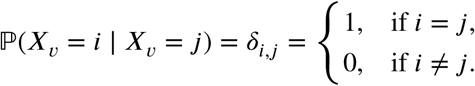
2. Going back in time, the first event is the admixture of populations 5 and 6 into 2. To process this event, we remove the set {2} and replace it with {5, 6}; the current collection of sets becomes {{1}, {3}, {5, 6}}. We make the new set {5, 6} the parent of the removed set {2} in 𝒯. In addition, we compute ℙ (*X*_2_ | *X*_5_ = *i, X*_6_ = *j*)]_*i,j*_, the conditional likelihood of the data beneath {5, 6} given the allele counts *X*_5_, *X*_6_. We obtain this by applying Lemmas 1 and 2 to the previous conditional likelihood ℙ(*X*_2_ | *X*_2_ = *i*)]_*i*_. Specifically, we apply Lemma 1 to “lift” the conditional likelihood at population 2 to the time point immediately below the admixture event; then, we apply Lemma 2 to obtain ℙ(*X*_2_ | *X*_5_, *X*_6_), the conditional likelihood at {5,6} immediately above the admixture event.
3. The next event has the alleles from 6 splitting off from population 3. To process this event, we merge the clusters containing the relevant populations ({5, 6} and {3}); then we remove 3,6 and replace them with their parent population 7, to obtain {5, 7} as the parent cluster of {5, 6} and {3} in. After this stage, our collection of sets becomes {{1}, {5, 7}}. To obtain ℙ(*X*_2_, *X*_3_ | *X*_5_, *X*_7_), the conditional likelihood of the data at the leaves beneath {5, 7}, we apply Lemma 3 to ℙ(*X*_3_ | *X*_3_) and ℙ(*X*_2_ | *X*_5_, *X*_6_). Lemma 3 computes the conditional likelihood at a split event when the children fall into separate clusters beneath the split (in this case, the child clusters are {3} and {5, 6}).
4. The next event is similar, with populations 1 and 5 merging into population 4. After combining the clusters {1} and {5, 7}, removing the merged populations 1 and 5, and adding in their parent 4, we are left with the collection {{4, 7}}. Similar to previous steps, we compute ℙ (*X*_1_, *X*_2_, *X*_3_ | *X*_4_, *X*_7_), the conditional likelihood of the data at the leaves beneath {4, 7}, by applying Lemma 1 to “lift” the conditional likelihoods ℙ(*X*_1_ | *X*_1_) and ℙ(*X*_2_, *X*_3_ | *X*_5_, *X*_7_) to the point immediately below the split event, and then apply Lemma 3 to compute the conditional likelihood at a split event above two independent clusters ({1}, {5, 7}).
5. The final event has populations 4 and 7 merging into population 8. We replace the cluster {4, 7} with its parent cluster {8} at the root of 𝒯. To compute ℙ(*X*_1_, *X*_2_, *X*_3_ | *X*_8_) from the child likelihoods ℙ(*X*_1_, *X*_2_, *X*_3_ | *X*_4_, *X*_7_), we first apply Lemma 1 to lift the likelihoods immediately below the split event, and then apply Lemma 4, which computes the conditional likelihood at the parent of a split when the children belong to the same cluster ({4, 7} in this case).

Finally, at each cluster {*v*_1_, *v*_2_,…} in 𝒯 with leaves {*l*_1_, *l*_2_,…}, we have computed 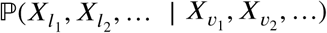, the conditional likelihoods at the leaves beneath {*v*_1_, *v*_2_,…}. But to apply equation (3), we ℙ((*X*_1_, *X*_2_, *X*_3_) = (1, 1, 0) | mutation at *v* with *X*_*v*_ = *i*). This is given by

ℙ((*X*_1_, *X*_2_, *X*_3_) = (1, 1, 0) | mutation at *v* with *X*_*v*_ = *i*)

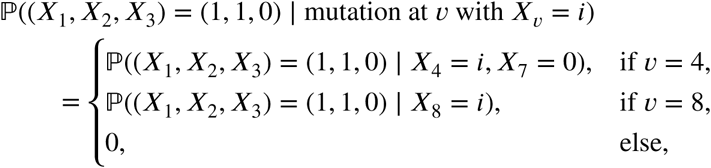

since we assume that each site experiences at most a single mutation in its history, and the only way derived alleles are observed in leaf populations 1,2 is if the corresponding mutation occurs in population 4 or 8.

#### Remark.

The observant reader may have noticed that in Figure 1, population 8 has only *n*_8_ = 5 alleles, despite its children having *n*_4_ + *n*_7_ = 7 > 5 alleles in total. This is due to the fact that there are only 5 alleles at the leaves, so there are at most 5 ancestors in any population at any point; thus, when populations 4 and 7 merge into population 8, we can drop two of the tracked lineages. To show this formally, we use a stochastic process called the *lookdown construction*, which is a version of the Moran model with a countably infinite number of lineages. However, this analysis requires a great deal of additional notation and is not essential to the remainder of the text, so we defer it to Appendix A.1.

### 3.2 Algorithms and formulas

We now describe the algorithm to construct the event tree 𝒯. We assume that the population graph 𝒢 has two types of topological events:

1. Population split: two child populations *u, v* split from each other; their parent population is *w*. Looking backward in time, *u, v* merge and become the population *w*.
2. Population admixture: a single child population *u*, inherits from exactly two parent populations *v, w*, with the probability that an allele comes from *v* (*w*, respectively) being *p* (1 − *p*, respectively).

Note that more complicated events, such as trifurcating splits or symmetric pulse migrations, may be expressed as a succession of these 2-way split and admixture events.

We provide pseudocode to construct the event tree 𝒯 in Algorithm 1. In words, we initially start with 𝒦 equal to a collection of singleton sets corresponding to the leaves of 𝒢. Processing each split or admixture event *E* back in time, we merge all blocks containing the child population(s) of *E*. Then we remove the child population(s) from this merged block, and add in the parent population(s). Within 𝒯, the new merged block is the parent of the blocks removed at this stage.

We now describe Algorithm 2, the dynamic program to compute the joint SFS. We need a bit more notation. For population *v*, let *n*_*v*_ be the number of samples with ancestry in *v*; we will be keeping track of *n*_*v*_ lineages within population *v*. Let *τ*_*v*_ denote the amount of time between the top and bottom events of *v*, and for let *Y* (*t*) denote the scaled population size of *v* at time *t* ∈ [0, *τ*_*v*_] above the base of *v*. Let 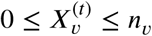 be the allele count within the *n*_*v*_ lineages of *v* at time *t* above its bottom, so 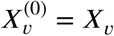 in our earlier notation.

At event *E* in 𝒯, let 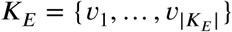 be the corresponding block of populations in𝒢. We define the conditional likelihood at *E* as

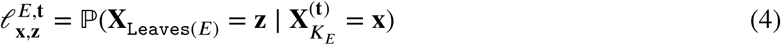

where 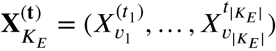 is the vector of allele counts in populations ***K***_*E*_ at times **t**, and **X**_Leaves(*E*)_ is the observed dataat the leaves beneath *E*.

In addition, we define the “partial SFS”

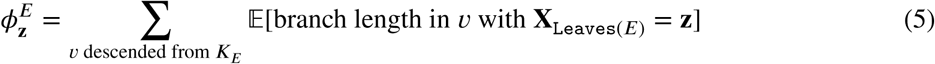

as the expected branch length at or below ***K***_*E*_ subtending **z** leaves. Note that 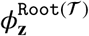 gives the desired final result, and corresponds to equation (3) in the previous subsection.

For the remainder of the subsection, we will fix **X** _Leaves(𝒢)_ = **z**, and drop the dependence on **z** in 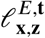 and 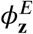. Algorithm 2 defines a dynamic program (DP) over the conditional likelihoods 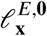 and partial SFS *Φ*^*E*^. The DP takes input vectors 𝓁^1^, …, *𝓁*^𝒟^ corresponding to the leaf populations Leaves 𝒢 = {1, …,𝒟}. If the inputs 𝓁^1^, …, *𝓁*^𝒟^ are set to indicator vectors corresponding to the observed counts (*X*_1_, …, *X*_𝒟_), the DP of Algorithm 2 will return the corresponding SFS entry, as stated in the theorem below:

#### Theorem 1.

*If* (*X*_1_, …, *X*_𝒟_) ≠ **O, n**, *then*

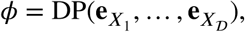

*where* 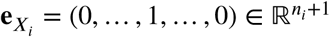 *denotes the vector with 1 at coordinate X*_*i*_ *and 0 elsewhere.*

We now present the formulas used by Algorithm 2, in a series of lemmas also used to prove Theorem 1. We start with a formula to “lift” 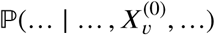 up to 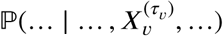. That is, this formula transforms a likelihood conditioned on 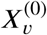, the allele count at the bottom of *v*, into a likelihood conditioned on 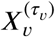,the allele count at the top of *v*.

#### Algorithm 1 Construct event tree 𝒯

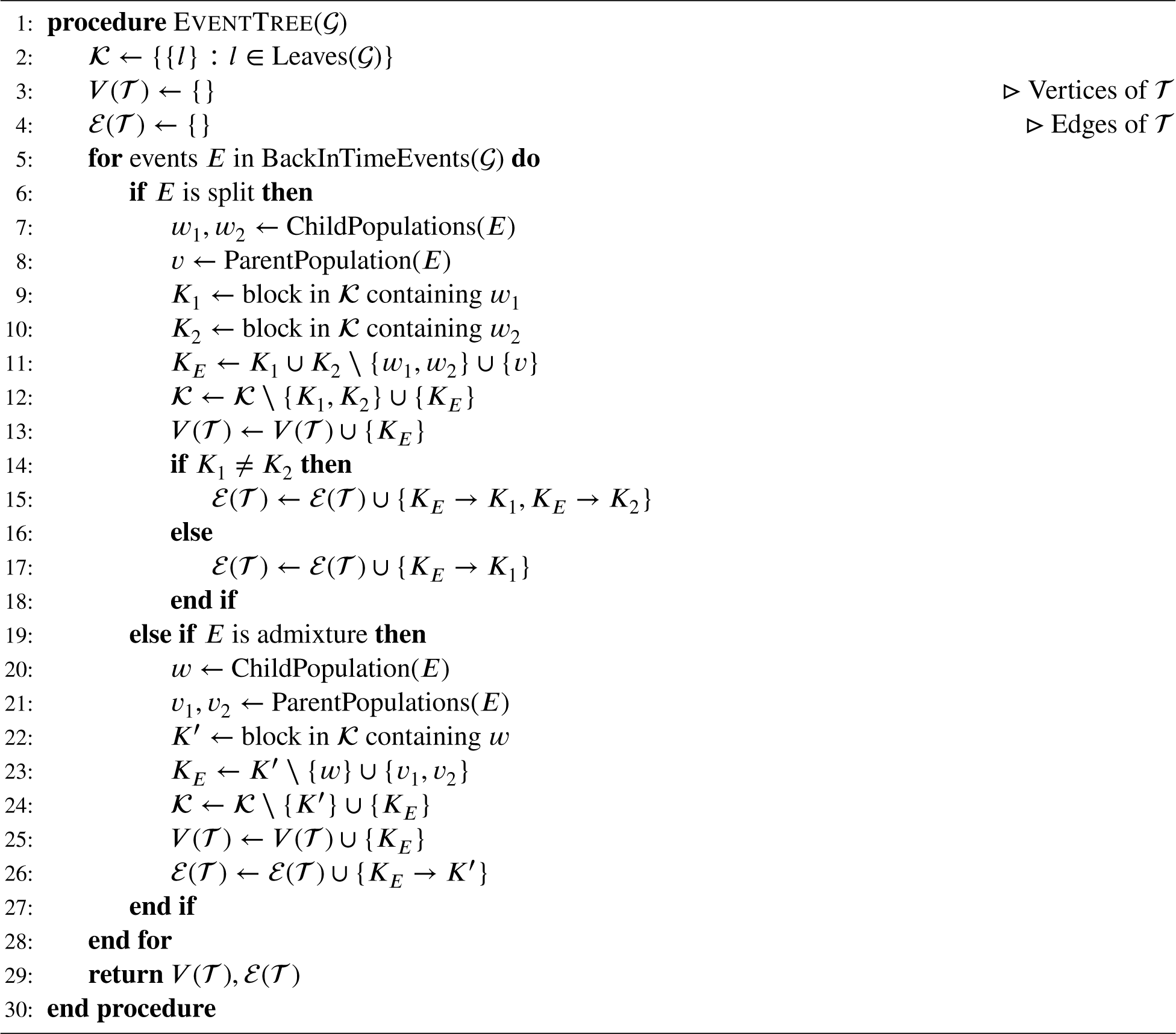

#### Lemma 1

(Lifting). *Let E be a split or admixture event with corresponding block K*_*E*_, *and let v* ∈ *K*_*E*_ *be a population within this block. Let* **x**_−*v*_ *a collection of allele counts on K*_*E*_ \ {*v*}, *and* **t**_−*v*_ *a collection of times on K*_*E*_ \ {*v*}. *Then, for* **x** = *k***e**_*v*_ + **x**_−*v*_ *and* **t** = *τ*_*v*_**e**_*v*_ + **t**_−*v*_, *the conditional likelihood of E is*

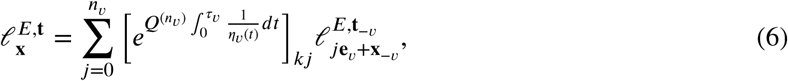

*where Q*^(*n*)^ ℝ ℛ ^(*n*+1)×(*n*+1)^ *is the transition rate matrix of the Moran model with n lineages; in particular*,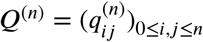 *and*

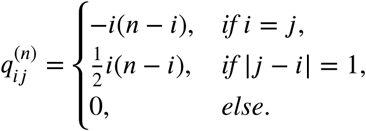

To process event *E*, we first use Lemma 1 to lift up the conditional likelihoods at the child populations, up to the time of *E*. We then apply one of Lemma 2, Lemma 3, or Lemma 4, to obtain the conditional likelihood at *E* from the lifted child likelihoods, depending on whether *E* is an admixture or split event, and whether the child populations of *E* fall into a single cluster or two independent clusters.

#### Algorithm 2 Dynamic program to compute the SFS *Φ*

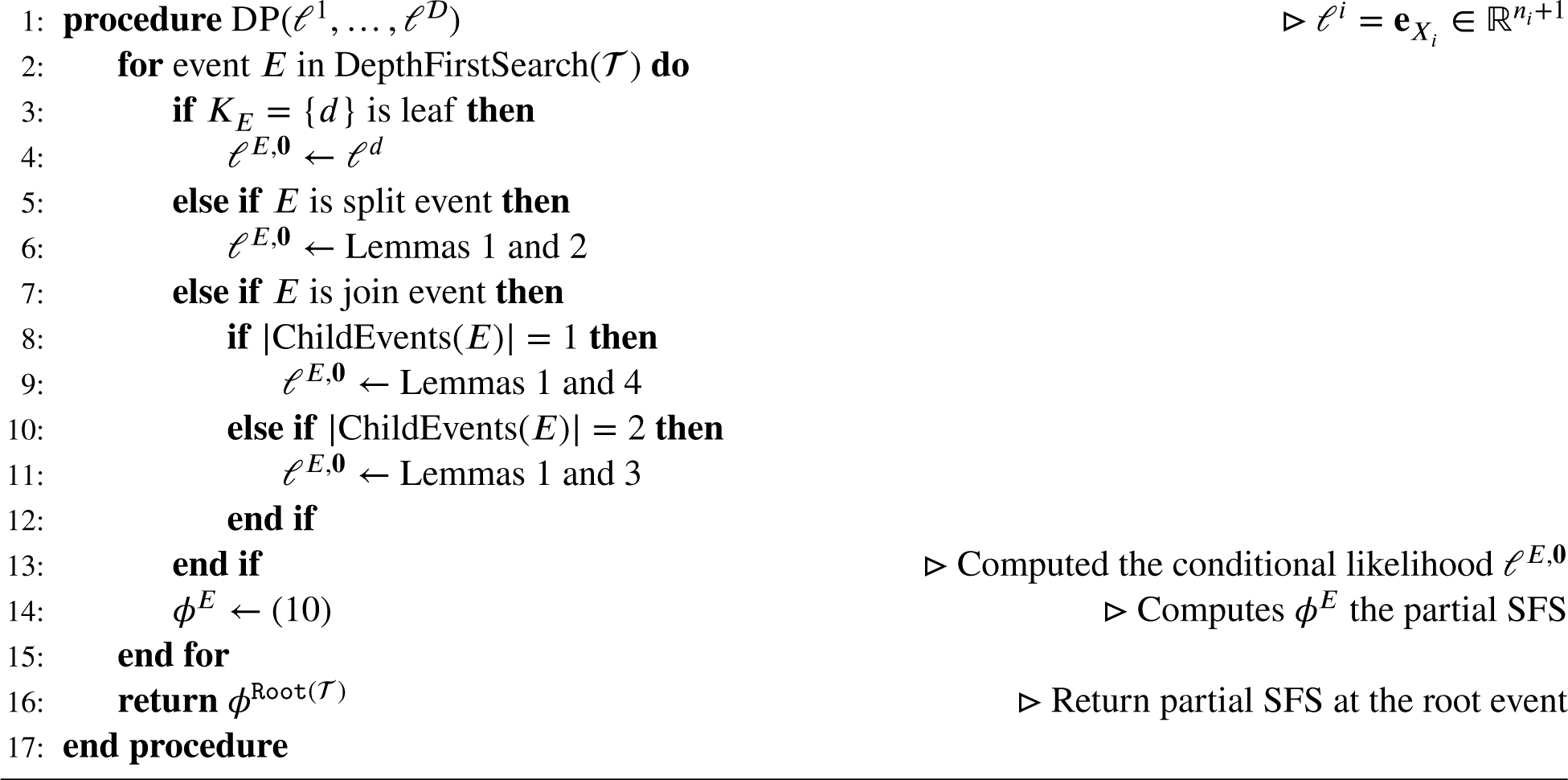

We first consider admixture events; Lemma 2 describes how to compute the conditional likelihood in this case. Let the child population be *w* and the parent populations be *v*_1_, *v*_2_. Each of the *n*_*w*_ lineages in *w* independently inherits from *v*_1_ with probability *q*_1_, or from *v*_2_ with probability *q*_2_ = 1 − *q*_1_. So the number of lineages inheriting from *v*_1_ is Binomial(*n*_*w*_, *q*_1_). Then, given that *m*_1_ alleles are inherited from *v*_1_ and *m*_2_ = *n*_*w*_ − *m*_1_ inherited from *v*_2_, the particular alleles inherited from *v*_1_ or *v*_2_ are chosen by sampling without replacement.

#### Lemma 2.

*(Admixture event) Let E be an admixture event, with child population w and parent populations v*_1_, *v*_2_. *Let E*^′^ *be the child event in 𝒯*. *Suppose each lineage in w comes from v*_1_ *with probability q*_1_, *and from v*_2_ *with probability q*_2_ = 1 − *q*_1_. *For K*_*E*_ *the population cluster at E, let* **x**_⋂_ *be a vector of allele counts on K*_*E*_ \ {*v*_1_, *v*_2_}. *Then the conditional likelihood of allele counts* 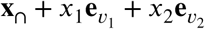 *at E is given by*

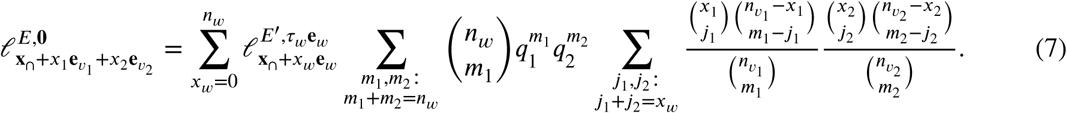

We next consider a split event *E*, with parent population *v* and child populations *w*_1_, *w*_2_. We first consider the case where *E* has 2 distinct children in *𝒯*, i.e., *w*_1_, *w*_2_ fall into 2 distinct blocks beneath *E*. Denote he corresponding child events as 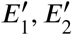 respectively Then the conditional likelihood at *E* is given by a convolution of the conditional likelihoods at 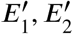, as described in Lemma 3:

#### Lemma 3.

*(Population split, 2 clusters) Let E be a split event with parent population v and child populations w*_1_, *w*_2_. *Suppose E has 2 child events* 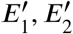, *with corresponding blocks* 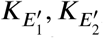, *where* 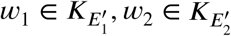. *Let* **x**_−1_ *be allele counts on* 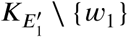 *and* **x**_−2_ *be allele counts on* 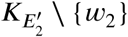. *Then the conditional likelihood at E is*

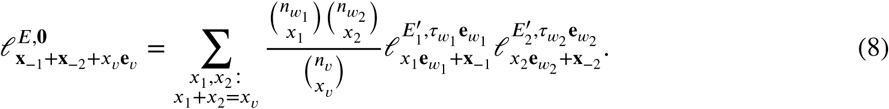

Now consider the case where the population split *E* has just 1 child event *E*′. That is, the child populations *w*_1_, *w*_2_ fall into the same cluster beneath. Then Lemma 4 describes how to obtain the conditional likelihood at *E* from the conditional likelihood at *E*′. This involves summing over the dimensions corresponding to *w*_1_, *w*_2_ within the conditional likelihood 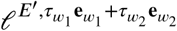. In addition, note that we may have 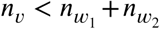 (recall *n*_v_ is the number of samples with ancestry in *v*). That is, after merging *w*_1_, *w*_2_ backwards in time, we may be keeping track of more alleles than originally sampled, allowing us to”drop” some extraneous non-ancestral lineages, as illustrated in the root population of Figure 1. This is done by multiplying the pseudoinverse of a matrix *B* whose entries are hypergeometric probabilities.

#### Lemma 4.

*(Population split, 1 cluster) Let E be a population split with exactly 1 child event E*′. *Denote the corresponding population clusters as K*_*E*_ *and K*_*E*_′. *Denote the parent population as v, the child populations as w*_1_, *w*_2_. *Let* **x**_n_ *be a vector of allele counts on the populations in K*_*E*_ ∩ *K*_*E*_′. *Define* 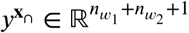 *and* 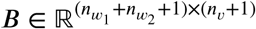 *to be the 0-indexed arrays with entries*

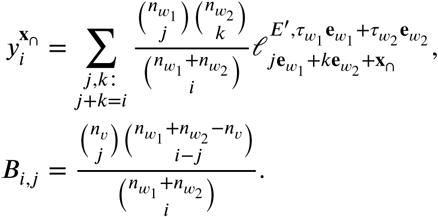

*Then the conditional likelihood at E is given by*

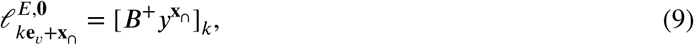

*with* ***B***^+^ *denoting the Moore-Penrose pseudoinverse of* ***B***.

Finally, having computed the conditional likelihoods at event *E*, we wish to compute the partial SFS *Φ*^*E*^ at the event. This is given by the recursive formula in Lemma 5, which involves the conditional likelihood at *E*, the expected number of mutations arising within each parent population, and the partial SFS at the child events.

#### Lemma 5.

*For an event E, let* 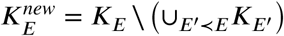 *be the populations newly formed at (i.e., formed by a population split or admixture at E). For* 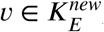, *let f*_*v*_ (*k*) *be the truncated SFS in population v*,

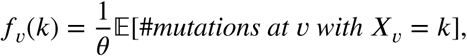

*which can be computed by the formulas in (Kamm et al., 2017). Then the partial SFS at E is given by*

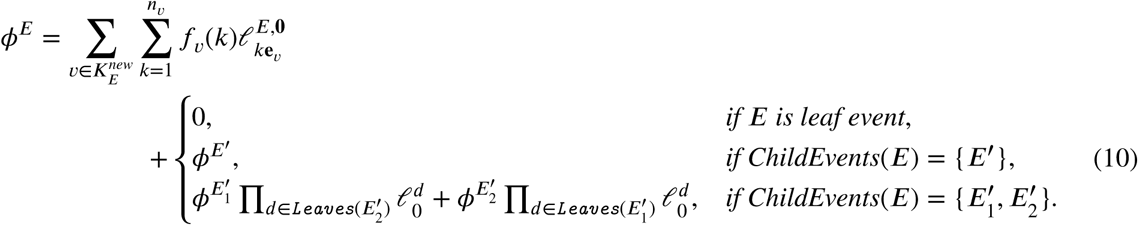

### 3.3 Normalizing constant and other linear functionals

To compute the probability 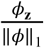 of a mutation having observed allele counts **z**, we need not just *Φ*_**z**_, but also the normalizing constant ‖*Φ*‖_1_ = Σ_**z**_*Φ*_**z**_ the expected total branch length.

Computing ‖*Φ*‖_1_ directly is in efficent because of the 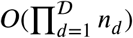 possible entries **z**. Instead, we can use Algorithm 2 to compute ‖*Φ*‖_1_, and many more statistics of the SFS, in the same time as a single entry:

#### Corollary 1.

*For* 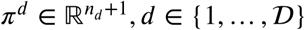, *the tensor dot product of the SFS Φ against* 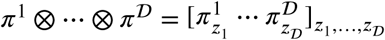 *is*

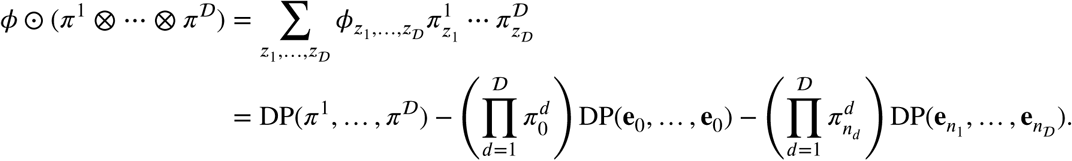

Corollary 1 says that for any rank-*K* tensor 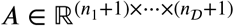 with 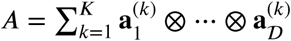,

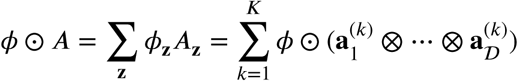

can be computed in *K* calls to DP(*π*^1^, …, *π*^𝒟^).^2^ In particular, the expected total branch length ‖*Φ*‖_1_ is given by

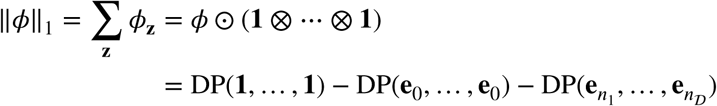

with **1** the vector with 1 at every coordinate.

Beyond the applications we explore here, we expect this result to be useful in a number of related settings. A number of population genetic statistics can be expressed as f ⊙ *A*, including Watterson’s estimator 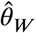 of the mutation rate (Watterson, 1975), Fay and Wu’s *H* statistic for positive selection (Fay and Wu, 2000), and Patterson’s *f*_2_, *f*_3_, *f*_4_ statistics for assessing topology (Patterson et al., 2012). Corollary 1 allows us to compute their expected values 𝔼[f ⊙ *A*] = *θΦ* ⊙*A*, and to construct test statistics from the deviance f ⊙ *A* − *θΦ* ⊙*A* under an appropriate null model.

An even wider class of population genetic statistics can be written as nonlinear functions of SFS-tensor products like 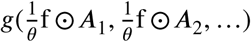; this class includes Tajima’s *D* statistic for selection (Tajima, 1989), the *F*_*ST*_ statistic for population structure (Holsinger and Weir, 2009), and Patterson’s *D* statistic for introgression (Patterson et al., 2012). These statistics may be viewed as plug-in estimators for g(*Φ* ⊙ A_1_, *Φ* ⊙ A_2_, …), which we can compute with Corollary 1. Note that these estimators are biased due to the nonlinear function *g*, but the bias can be estimated via block jackknife, and will typically be small since 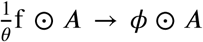 almost surely as the number of independent SNPs grows.

Another interesting linear statistic of the SFS that can be computed with Corollary 1 is 𝔼*T*_MRCA_], the expected time of the most recent common ancestor. In particular, let *d* be any leaf population; for simplicity assume *d* is sampled at the present (i.e. *d* is not archaic). Then

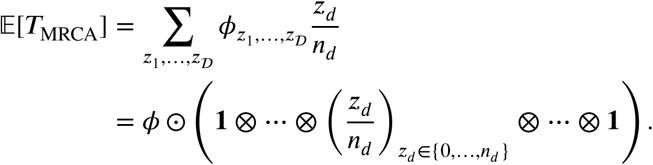

To see this, note that *T*_MRCA_ is proportional to the expected number of mutations hitting an arbitrary lineage in *d*, and if a mutation has configuration **z** = (*z*_1_, …, *z*_*𝒟*_) derived copies, then the chance of hitting the lineage is 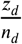 (Zeng et al., 2006).

## 4 Application

We tested our method on a demographic inference problem in human genetics that is currently of interest. Lazaridis et al. (2014) showed that genetic variation in present-day Europeans suggests an admixture model involving three ancestral meta-populations: Ancient North Eurasian (ANE), Western Hunter Gatherers (WHG), and Early European Farmers (EEF). They also showed that EEF contains ancestry from a source that is an outgroup to all non-African populations, and yet shares much of the genetic drift common to non-African populations; they dubbed this ancestry component as “Basal Eurasian” ancestry. Later work (Lazaridis et al., 2016) showed that Basal Eurasian ancestry is shared by ancient and contemporary Middle Eastern populations, and is correlated with a decrease in Neanderthal ancestry, implying that Basal Eurasian ancestry contains lower levels of Neanderthal admixture when compared with non-Basal ancestry. The results from Lazaridis et al. (2014, 2016) were based on several related methods for modeling covariances in population allele frequencies, most notably qpGraph and qpAdm (Patterson et al., 2012; Haak et al., 2015). These methods are computationally efficient and robust, but are unable to infer the timing of demographic events.

We applied `momi2` to estimate the strength and timing of basal Eurasian admixture into early European farmers, and the split time of the basal Eurasian lineage. To do this, we built a demographic model relating 12 samples from 8 populations, shown in Figure 2. These samples consisted of the Altai Neanderthal (Prüfer et al., 2014); the 45,000 year old Ust’Ishim man from Siberia (Fu et al., 2014); 3 present-day populations (Mbuti, Sardinian, Han) with 3, 2, and 2 samples respectively; and 3 ancient samples representing the European ancestry components identified by Lazaridis et al. (2014): a 7,500 year old sample from the Linearbandkeramik (LBK) culture (representing EEF), an 8,000 year old sample from the Loschbour rock shelter in Luxembourg (representing WHG), and the 24,000 year old Mal’ta boy (“MA1”) from Siberia (representing ANE). After data cleaning, our dataset consisted of 2.4 × 10^6^ autosomal transversion SNPs. See Appendix A.3 for more details about the data.

**Figure 2:**
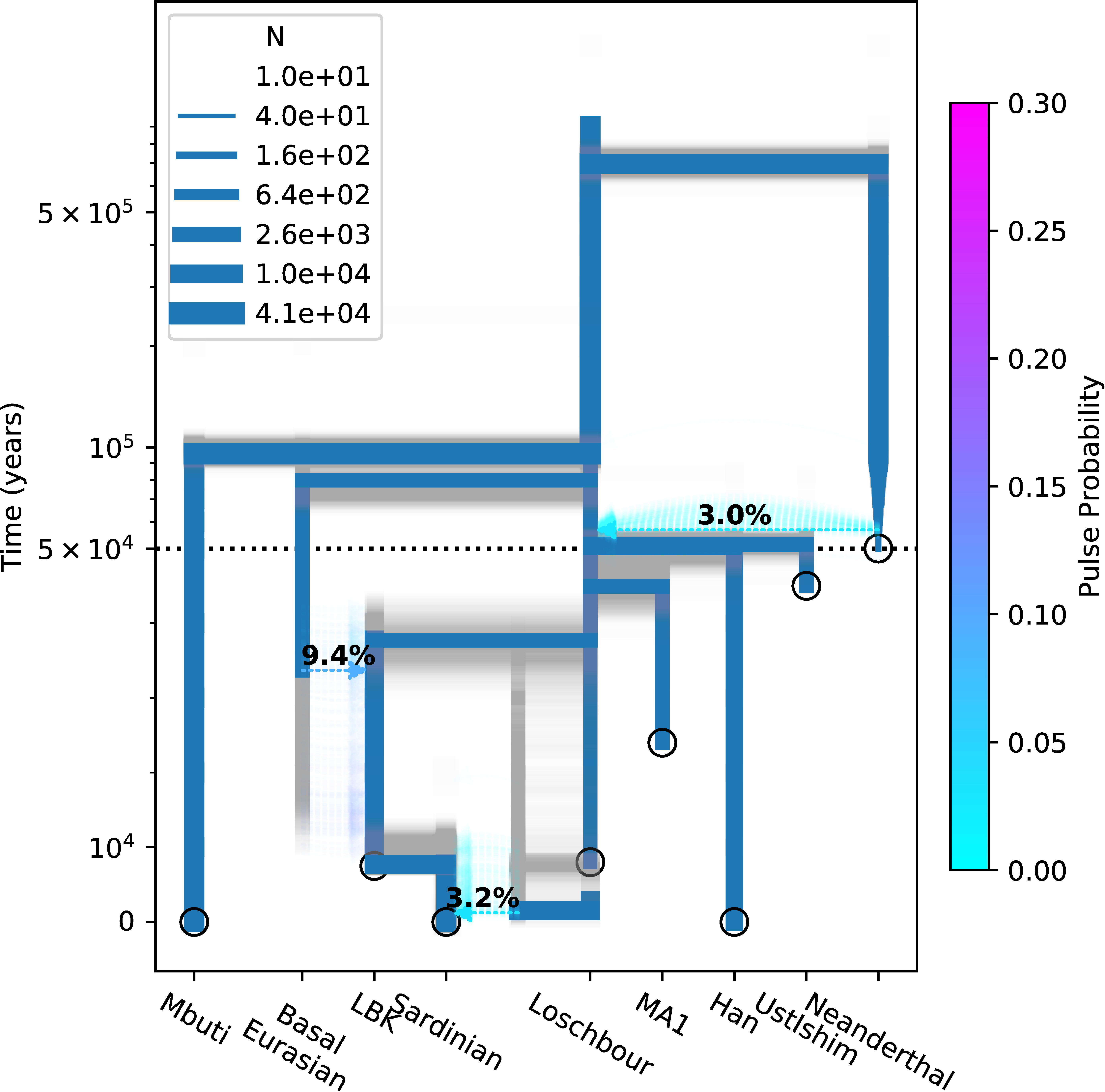
Inferred model and bootstraps for the 11 population demography described in Section 4. In the foreground (blue) is our point estimate from maximum composite likelihood; in the background (gray) are 300 bootstrap re estimates, which were created by splitting the data into 100 equally sized contiguous blocks, resampling these blocks with replacement, and refitting the model. The y-axis is linear below 5 × 10^4^, then follows a logarithmic scale above 5 × 10^4^.

To construct the topology of the model in Figure 2, we first obtained a tree by neighbor joining (Saitou and Nei, 1987), then added 3 extra admixture events reflecting prior knowledge, as well as a recent Neanderthal population decline starting at the Mbuti-Eurasian split. We inferred split times, population sizes (including the Neanderthal decline), and admixture times and proportions by maximizing a composite likelihood, given by the product of the likelihoods at every SNP:

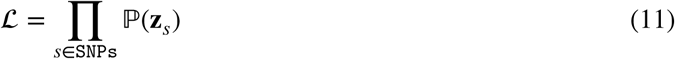

where 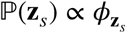 and was computed by `momi2`. The low-coverage of the MA1 sample and the deep divergence of the Neanderthal sample may cause bias in SFS entries where only these samples contain derived alleles; we thus excluded these entries and corrected the normalizing constant appropriately (see Appendix A.3).

We used automatic differentiation to compute the gradient ∇ logℒ, which we used to search for the optimum of ℒ. We constructed nonparametric bootstrap confidence intervals by splitting the genome into 100 equally sized blocks, resampling these blocks to create 300 bootstrap datasets, and re-inferring the demography for each bootstrap dataset. We also used 300 parametric bootstraps to assess how well we could infer the demography under simulated data; for each parametric bootstrap dataset, we used `msprime` (Kelle-her et al., 2016) to simulate ten 300 Mb chromosomes from our initial point estimate, and re-inferred the demography. Note the nonparametric bootstrap is better able to account for model misspecification, and we use it for all confidence intervals reported below.

Our inferred demography, along with nonparametric bootstrap re-estimates, are shown in Figure 2 and Table 2. Our parametric bootstrap estimates are shown in Figure 3. We inferred a pulse of 0.094 (95% CI of 0.049-0.174) from the ghost Basal Eurasian population to EEF ancestry (LBK), substantially less than the 0.44 inferred by (Lazaridis et al., 2014). This admixture was inferred to occur 33.7 kya (95% CI of 10.8-41.1 kya), shortly after the Loschbour-LBK split at 37.7 kya (95% CI of 32.2-42.3 kya). The split time of the ghost Basal Eurasian lineage from other Eurasians was inferred at 79.8 kya (95% CI of 67.4-101 kya). Other parameters were broadly in line with previous estimates, such as a Mbuti-Eurasian split of 96 kya, a Han-European split of 50 kya, a Neanderthal split of 696 kya, and Eurasians deriving 0.03 of their ancestry from Neanderthal (Terhorst et al., 2017; Green et al., 2010; Meyer et al., 2016).

**Table 2:**
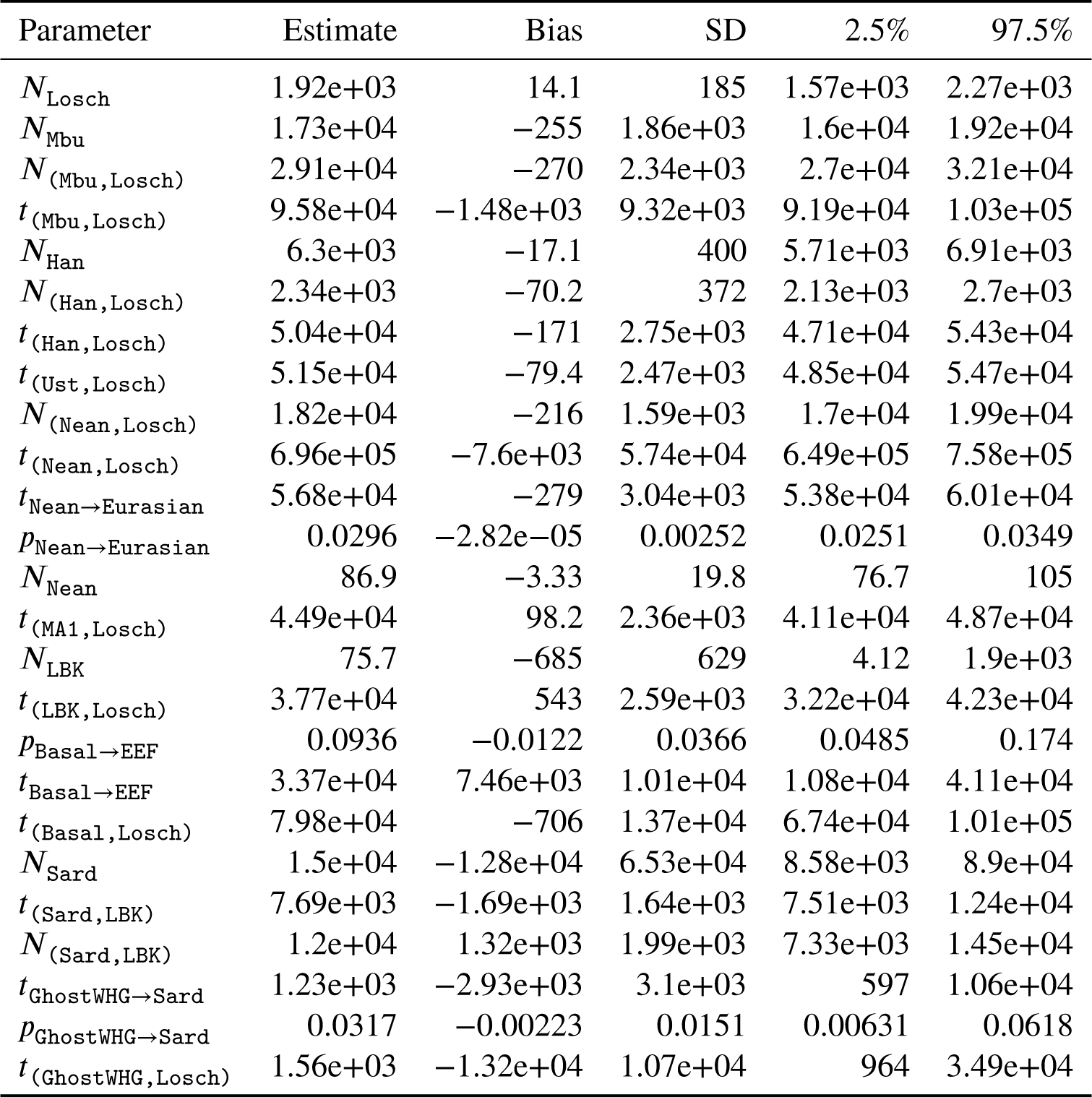
Estimated parameters of the demography in Figure 2, along with nonparametric bootstrap estimates of the bias and standard deviation, and 95% bootstrap quantiles. We use (A,B) to denote the ancestor of A and B; *t*_*v*_ and *N*_*v*_ to denote the height and size at vertex *v*; and *t*_A→B_ and *p*_A→B_ to denote respectively the time and strength of an admixture arrow from A to B.

**Figure 3:**
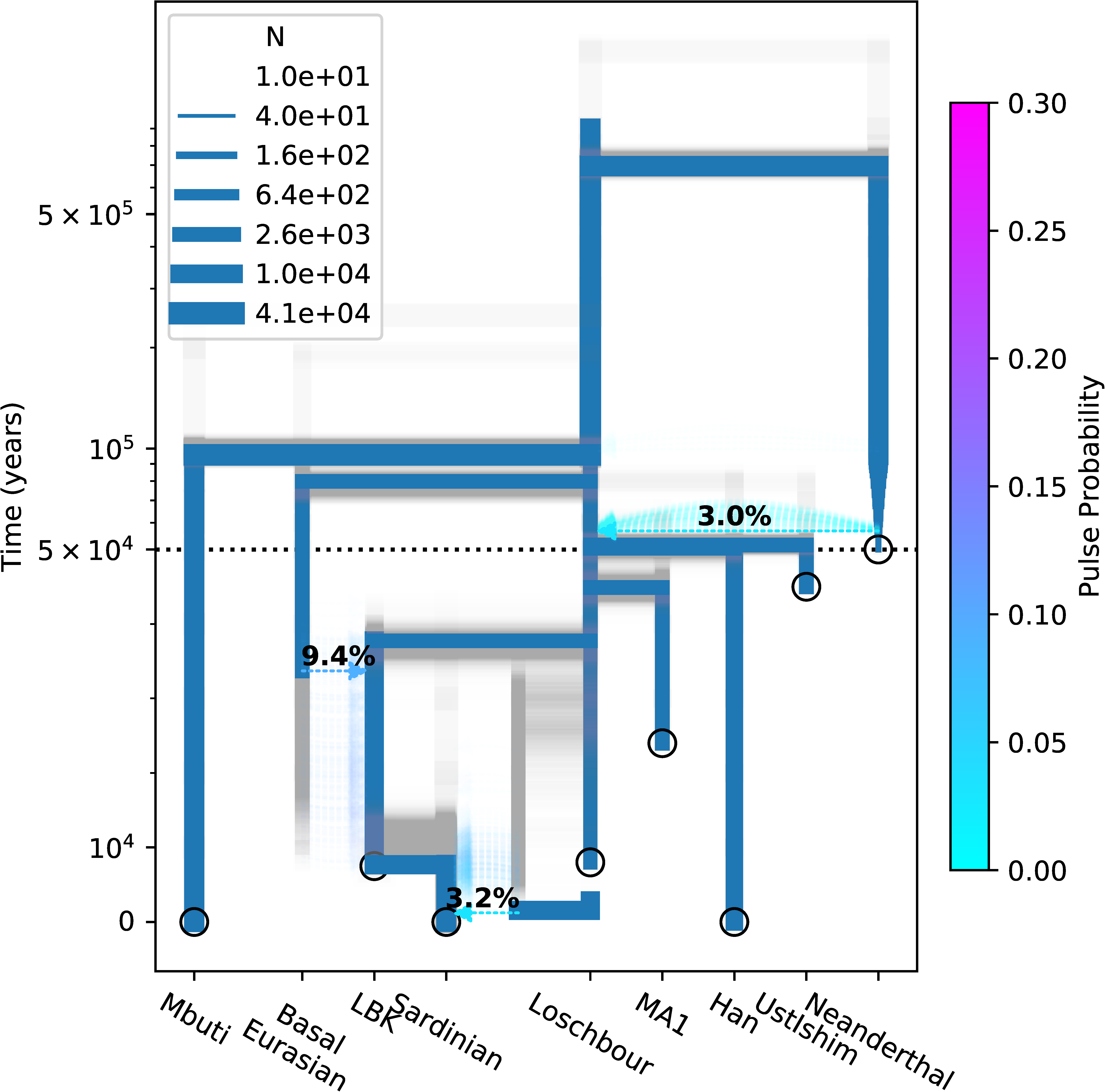
Parametric bootstraps for the demography inferred in Figure 2. The inferred demography (blue tree) was used to simulate 300 bootstrap replicates. Re-running our method on each replicate produced a new inferred demography which is plotted in gray.

Inferring the optimal demography from start to finish took 2.5 hours on a laptop with 4 CPU cores, and used 2 GB RAM. The 300 bootstraps were run separately on a high-performance compute cluster. To our knowledge, no other method can infer this demographic model using the full SFS. The `moments` software package (Jouganous et al., 2017) is capable of computing the SFS for up to 5 populations, less than the 8 populations here, though it can scale to more individuals per population than `momi2`. While the `fastsimcoal2` software package (Excoffier et al., 2013) is capable of handling demographies of this size and larger, it doesn’t compute the full, exact SFS, and also does not include an option for the ascertainment scheme we use here (excluding mutations private to Neanderthal and MA1).

We also used our model to produce estimates of the human mutation rate, which we estimated as 1.22×10^−8^ per base per generation, with a bootstrap-quantile 95% CI of (1.12, 1.32) × 10^−8^, closely matching previous estimates of the human mutation rate (Scally, 2016). To obtain this estimate, we compared the observed nucleotide diversity with that expected under the inferred demography, adjusting by the empirical transition to transversion ratio (Appendix A.3). Estimating the mutation rate was possible here because we did not use a prespecified mutation rate to estimate the model in Figure 2, instead using the known ages of the Ust’Ishim, LBK, Loschbour, and MA1 samples to calibrate dates.

## A Appendix

### A.1 Model and notation

In this section, we formally define the stochastic process underlying our model, and introduce some additional notation needed for the proofs in Section A.2.

The main stochastic process we use is the *lookdown construction* of the Moran model (Donnelly and Kurtz, 1996; Donnelly et al., 1999), a variant of the Moran model where copying only occurs in one “direction” (as in Figure 4). We will make use of certain conditional independence properties that result from this one-way copying. However, we note that there is a simple coupling between the lookdown and standard Moran models, and these two models generate data with the same distribution.

**Figure 4:**
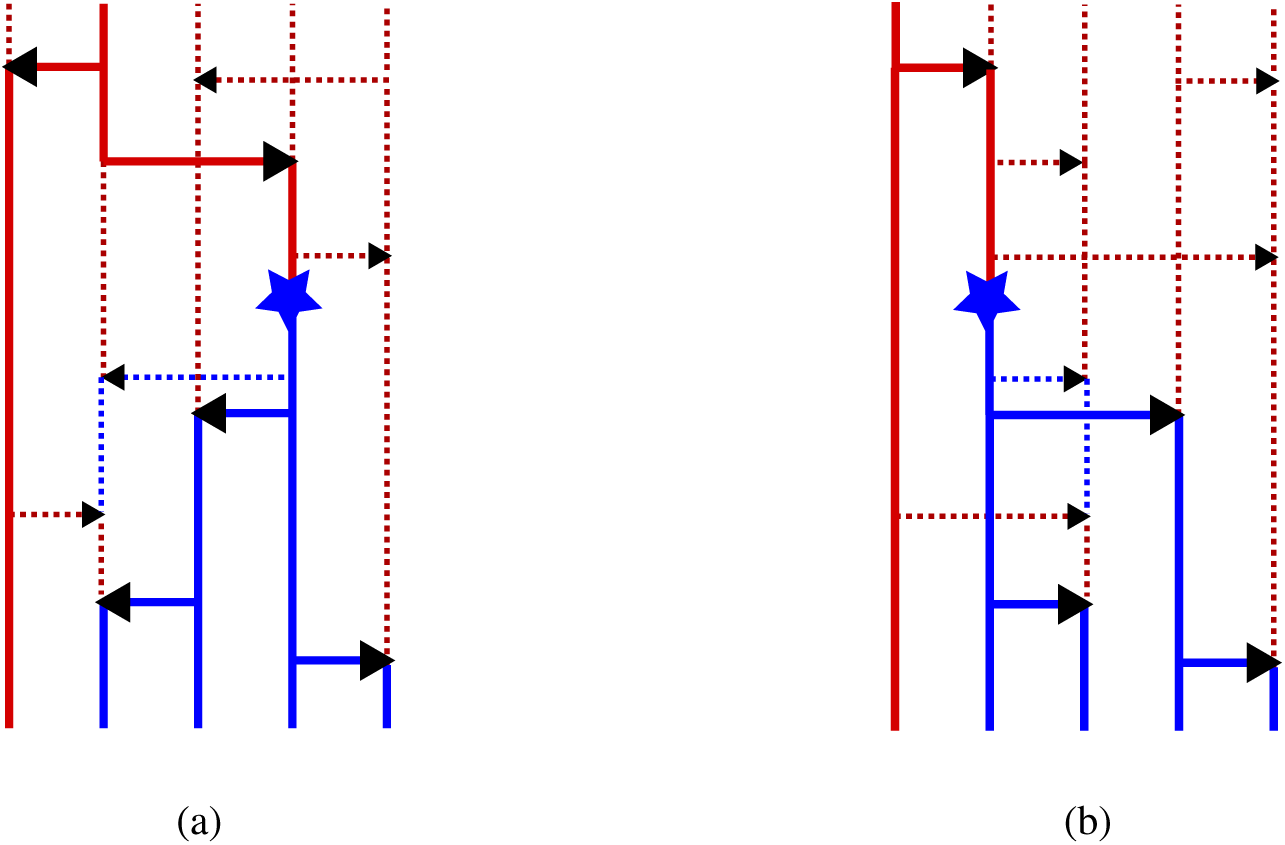
Standard and lookdown Moran models. (a) illustrates the standard Moran model, with copying in both directions; (b) illustrates the lookdown Moran model, with copying always from left to right. The sample genealogy (i.e., the lineages ancestral to present day samples) is in solid. The law of the genealogy is the coalescent under both models, and so the present-day samples of the two models are equal in distribution.

We now describe our lookdown model in more detail. Within each of the leaf populations we consider a countably infinite number of lineages, each with a unique label in Z_+_. We arbitrarily assign the (*n*_1_, …, *n*_𝒟_) sampled lineages to the lowest labels {1, 2, …, *n*_tot_}, where 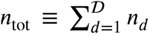, and arbitrarily assign the remaining unsampled lineages to labels {*n*_tot_ + 1, *n*_tot_ + 2, …}. Each lineage extends infinitely backwards in time, and at each admixture event, each lineage randomly chooses the parent population it extends into. In addition, each lineage has an allele, which changes through time due to mutation and copying events.

Similar to the usual Moran model, copying between a pair of lineages in population *v* occurs at rate 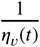, where *η*_*v*_ (*t*) is the scaled population size of *v*. However, copying only occurs in one direction, from lineages with lower labels to lineages with higher labels (Figure 4). So if two lineages are labeled *i* and *j* respectively, with *i < j*, then the copying event *i* → *j* occurs at rate 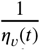, whereas the reverse copying event *i* +- *j* never happens (has rate 0). By contrast, in the standard (non-lookdown) Moran model, both *i* → *j* and *i* +- *j* copying events happen at rate 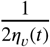. However, under both models, two lineages going backwards in time coalesce at rate 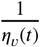, as in the coalescent. Thus, the sample genealogies under both models follow the same multipopulation coalescent distribution.

Denote the labeled lineages in population *v*, and their alleles at height *t*, by

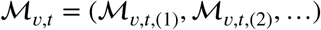

where ℳ_*v,t*,(*i*)_ = (`label`_*v*,(*i*)_, `allele`_*v,t*,(*i*)_) ∈ ℤ_+_ × {0, 1} is a pair, consisting of the *i*th lowest label at *v*,along with its corresponding allele at height *t*, ∈ [0, *τ*_*v*_). Note that we have ordered the lineages by their labels, so that `label`_*v*,(*i*)_,< `label`_*v*,(*i*+1)_ Also note that we measure the time *t* from the bottom of *v*, so that allele*v*,0,(*i*) denotes the *i*th allele at the bottom of *v*, and allele*v,τ*,(*i*) the *i*th allele at the top of *v*.

Let 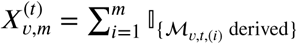 denote the number of derived alleles among the lowest *m* lineages at *v, t*. Let *n*_*v*_ = Σ_*d⪰v*_ *n*_*d*_ be the number of samples in leaves below *v*. During computation, we will only need to keep track of the lowest *n*_*v*_ lineages in *v*, so we denote 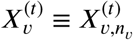 to be the number of derived alleles among the first *n*_*v*_ lineages at. *v, t.*

Finally, for an event *E* with *K*_*E*_ = (*v*_1_, …, *v*_*k*_) the corresponding block of populations in Algorithm 1, let **t** = (*t*_1_, …, *t*_*k*_) be a corresponding set of times within each population *v*_1_, …, *v*_*k*_. We define 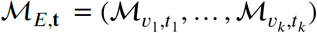 as the labeled alleles at each of (*v*_1_, *t*_*k*_), …, (*v*_1_, *t*_*k*_).

### A.2 Proofs

The following two lemmas will be useful for several of the proofs below:

#### Lemma 6.

*The distribution of* M_*E*,**t**_ *is invariant to finite permutations of the labels within any population v* ∈ *K*_*E*_. *Furthermore, the labels are independent of the alleles.*

*Proof.* By construction, none of the lineages within the populations in *K*_*E*_ are ancestral to each other. Thus the sample genealogy of any finite subsample of the lineages is the multipopulation coalescent, because going backwards in time, coalescence (copying) between each pair of lineages occurs at rate 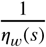 at vertex *w* time *s*. The invariance to permutation, and independence of alleles and labels, follows from the coalescent.

#### Lemma 7.

*For event E and population u* ∈ *K*, *let* ℐ *c* {*n* + 1, *n* + 2, …} *be a (possibly random) collection of integers greater than n*_*u*_, *and let* 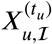 *be the number of derived lineages in* 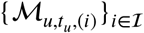. *Then* **X**_*Leaves*(*E*)_ *is conditionally independent of* 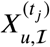 *the allele counts on ℐ, given* 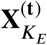 *the allele counts on the first* 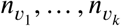 *lineages of K* = {*v*_1_, …, *v*_*k*_}.

*Proof.* Integrate over 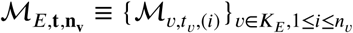 the first 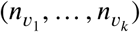 labeled alleles within *K*_*E*_, and 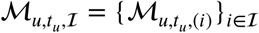 the labeled alleles at ℐ:

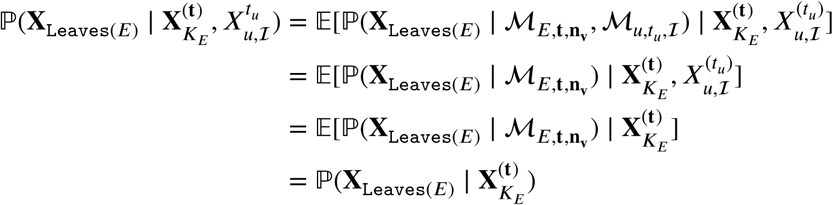

with the second equality because higher lineages cannot copy to lower lineages (so 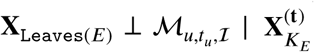), and the third equality because of the exchangeability and independence from Lemma 6 (so given 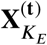, the alleles of 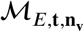 are ordered by a uniform permutation independent of 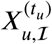, and the labels of 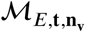 are independent of 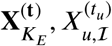.

#### A.2.1 Proof of Theorem 1

Lemmas 2,4,3 give formulas for the partial likelihood 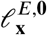 using terms like 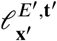, where *E*′ ∈ `Children`(*E*) and *E* is an admixture, 1-cluster split, or 2-cluster split, respectively. The terms 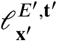 in turn can be computed from the partial likelihoods 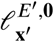 using Lemma 1; then Lemma 5 provides a formula for *Φ*^*E*^ in terms of 𝓁^*E*,**0**^ and the partial SFS 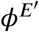.

Algorithm 2 traverses the event tree 𝒯 in a depth-first search, applying Lemmas 5,1,2,4,3 to compute 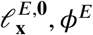, *Φ*^*E*^ at each event *E* from their values at the children of *E*. The input to Algorithm 2 are the likelihoods 𝓁^{*d*},**0**^ at the leaves, and the output is the partial SFS *Φ*^Root(𝒯)^ at the root.

Thus, since

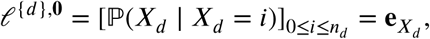

it follows that DP 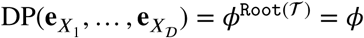.

##### A2.2 Proof of Lemma 5

Without loss of generality assume *0* = 1. Then *Φ^E^* is the expected number of mutations at or below *K*_*E*_ with observed counts **X**_Leaves(*E*)_. We split *Φ*^*E*^ into two parts: the mutations within 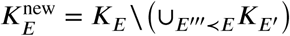 (i.e. the populations formed by a split or join at *E*); and the mutations that occur in ∪_*E′≺E*_ *K*_*E′*_, the populations that arise strictly below *E*.

The first part, of mutations at the new populations of *E*, is given by

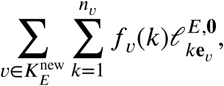

since *f*_*v*_(*k*) is the expected number of mutations arising at *v* with *X*_*v*_ = *k*, and 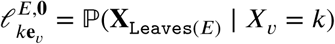.

For the second part, of mutations strictly below *E*, we split into three cases: either *E* is a leaf event, or *E* has a single child event, or *E* has two child events. In the first case, if *E* is a leaf event, then no mutations can occur below *E*. In the second case, if *E* has a single child event *E*′, then the expected number of mutations strictly below *E* is simply 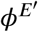 by definition.

Finally, if *E* has two child events 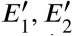, then 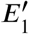 and 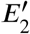 share no leaves, so a mutation underneath 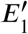 will have no derived alleles in `Leaves` 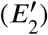 and vice^1^versa. Thus, the number of mutations strictly below *E* is

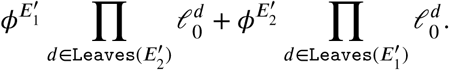

##### A.2.3 Proof of Lemma 1

Define a “quasi-lookdown” Moran model ℳ*, which is identical to ℳ, except within the *n*_*v*_ lowest lineages of *v*, where we allow copying in both directions at rate 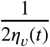 (as in the non-lookdown Moran model).

For an event *E* with populations *K*_*E*_ = (*v*_1_, *v*_2_, …) and corresponding times 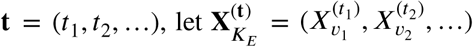 be the corresponding allele counts. Next, define 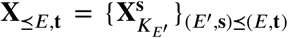 as the sample path of allele counts below *E*, **t**, where (*E*′, **s**) :s (*E*, **t**) if either *E* is above, or *E* = *E*′ and **s ≤ t** component-wise. It will suffice to show 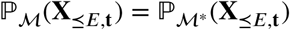, because then for **t** = *τ*_*v*_ **e**_*v*_ + **t**_−*v*_,

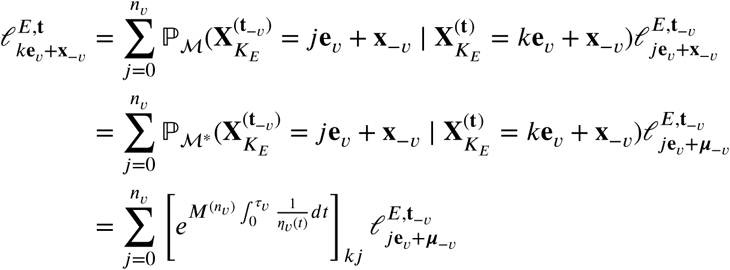

as desired.

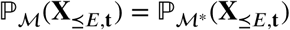 follows from a coupling argument. Let 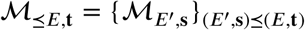 the sample path of ℳ below *E*, **t**. We can map the partial sample paths of 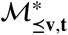 onto those of ℳ_s⪯**vt**_ as follows: moving from the bottom to the top of *v*, whenever a lower label is copied over by a higher label, swap the labels of the lineages above the copying. Then the relabeled sample path has the same distribution as the lookdown construction, since the allele with the higher label is always copied over, and the rate of copying between pairs of lineages is 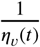. Since this relabeling also leaves 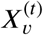 unchanged, we have 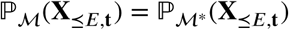.

##### A.2.4 Proof of Lemma 2

First note that **X**_Leaves(*E*)_ = **X**_Leaves(*E′*)_ and

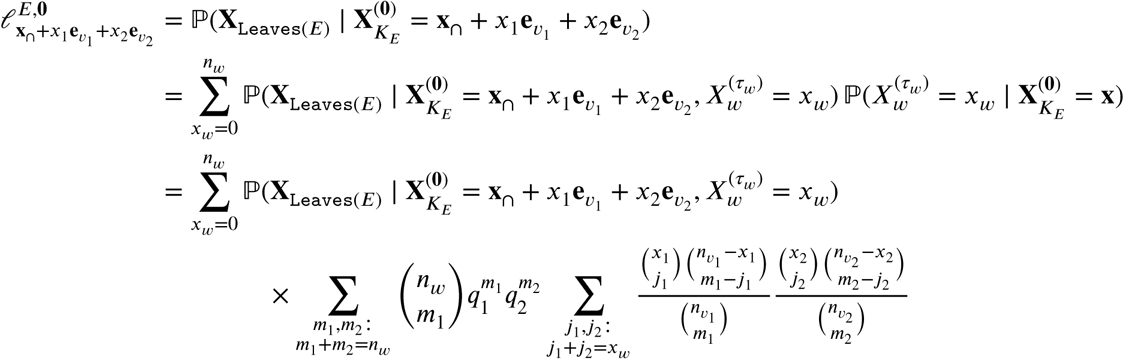

by sampling *n*_*w*_ alleles in *w* from 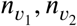 alleles in *v*_1_, *v*_2_, which are exchangeable by Lemma 6.

Next, consider the *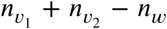* highest alleles of *v*_1_, *v*_2_, and let ℐ ⊂ {*n*_*w*_ + 1, *n*_*w*_ + 2, …} be their relative positions within *w*. Then we conclude the proof by noting that **X**_Leaves(*E*)_ = **X**_Leaves(*E′*)_ and applying Lemma 7, to get

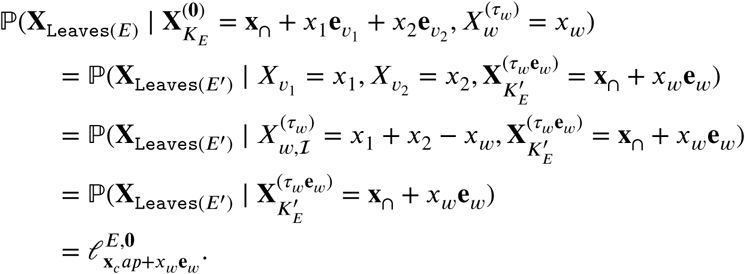

##### A.2.5 Proof of Lemma 4

Let 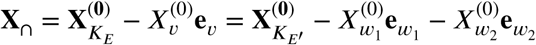 be the vector of allele counts on *K*_*E*_ ∩ *K*_*E′*_ at times **0**.Then note that

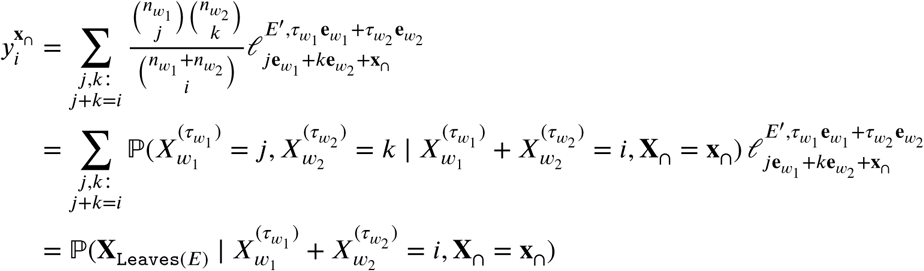

with the second equality following from exchangeability (Lemma 6) and the third equality from **X**_Leaves(*E*)_ = **X**_Leaves(*E′*)_.

Next note that

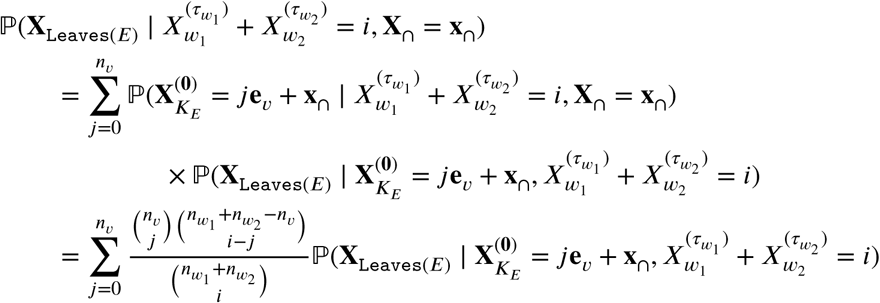

with the second equality again due to exchangeability (Lemma 6).

Define ℐ ⊂ {*n*_*v*_ + 1, *n*_*v*_ + 2, …} so that {1, …, *n*_*v*_} ⋃ ℐ are the indices in *v* of the first 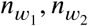 alleles in *w*_1_, *w*_2_. Then because *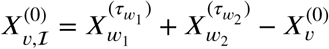* and Lemma 7,

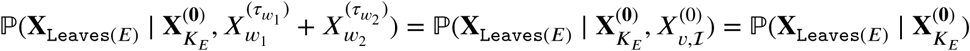

and thus

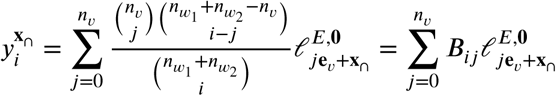

so letting 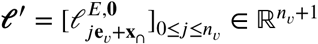, we have 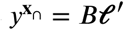 and therefore 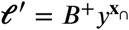.

##### A.2.6 Proof of Lemma 3

Notice that

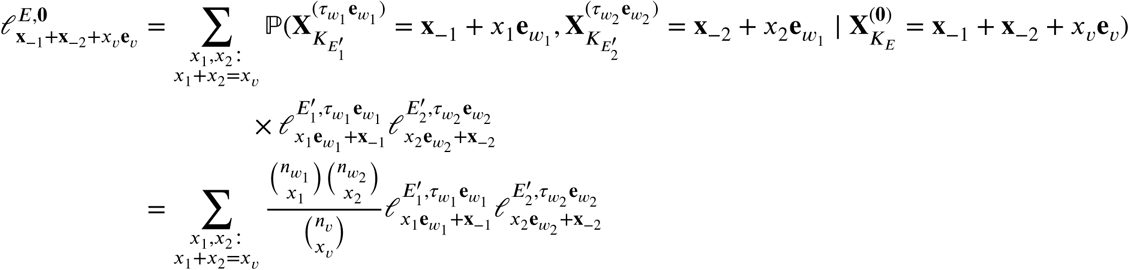

with the first equality from the Markov property of the Moran process, and the second equality following from the exchangeability of the *n*_*v*_ alleles at vertex *v* (Lemma 6).

##### A.2.7 Proof of Corollary 1

Below, we will prove DP(𝓁^1^, …, 𝓁^*𝒟*^) is a multilinear function of the input vectors 𝓁^1^, …, 𝓁^*𝒟*^. The result immediately follows from this, because then

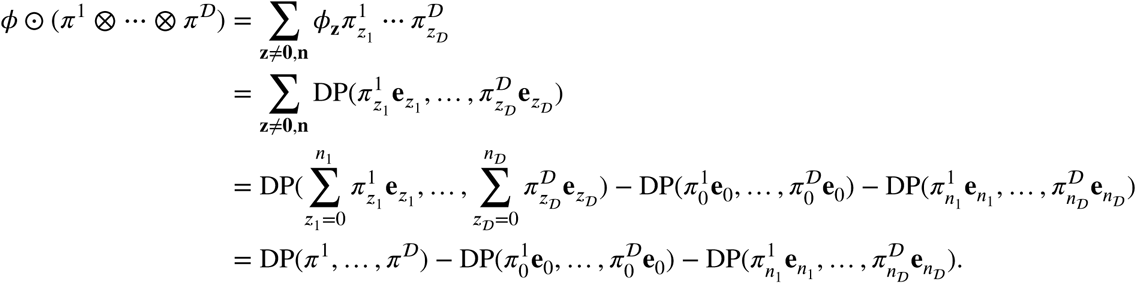

We now show DP(𝓁^1^, …, 𝓁^*𝒟*^) is a multilinear function of 𝓁^1^, …, 𝓁^*𝒟*^. We start by showing that if event *E* has leaf populations `Leaves`(***E***) = (*d*_1_, …, *d*_*L*_), then 𝓁^*E*,**t**^ is a multilinear functions of 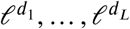. We show this by induction over (***E*, t**). The base case, where **t** = 0 and ***E*** is a leaf with ***K***_*E*_ = {*d*_*i*_}, is trivially true because 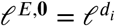.

For the next case, we note that (6), (7), and (9) express 𝓁^*E*,t^ as a tensor product of *l*^*E*′,*t*′^ with a matrix, where (*E*′, **t**′) ≺ (*E*, **t**) and `Leaves`(*E*) = `Leaves` (*E*′). *l*^*E*′,*t*′^ is a multilinear function of 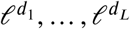 by induction, hence 𝓁^*E*,**t**^ is also.

Similarly, (8) expresses 𝓁^*E*,0^ as a product of 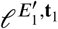 and 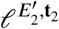 where 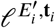 is a multilinear function of 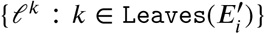 by induction, and furthermore 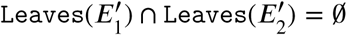 and 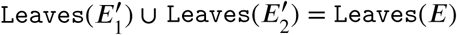 Thus 𝓁^*E*,**0**^ is a multilinear function of 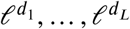.

Thus, each of the operations (6), (7), (8), (9) in Algorithm 2 preserves multilinearity of 𝓁^*E*,**t**^, and thus each 𝓁^*E*,**t**^ is a multilinear function of its leaf vectors by induction. Finally, a similar induction argument shows that each *ϕ*^*E*^ is a multilinear function of {𝓁^*k*^ : *k* ∈ `Leaves`(*E*)}, since (10) expresses *ϕ*^*E*^ as a sum of multilinear functions by induction.

### A.3 Application supplement

#### A.3.1 Data

Table 3 gives the populations and samples we used for our example application in Section 4. All samples except for MA1 were taken from the SGDP dataset (Mallick et al., 2016). We added on the additional low-coverage MA1 sample (Raghavan et al., 2014) to represent the Ancient North Eurasian (ANE) component of European ancestry. Due to the low-coverage of MA1, we only sampled a single random allele from it at each site, using a GATK pileup (McKenna et al., 2010) and restricting ourselves to reads with quality ≥ 30.

**Table 3:**
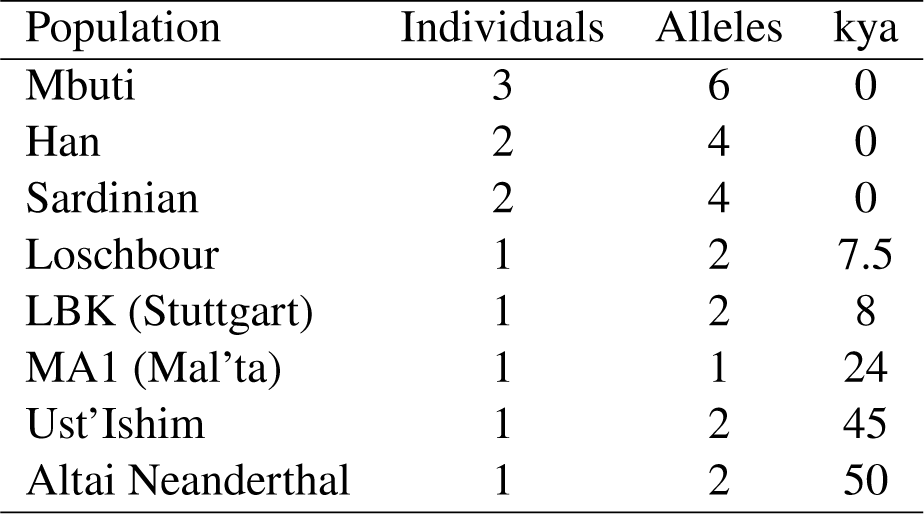
Populations and samples used for the example application in Section 4. We only used 1 random allele for MA1 due to low coverage. The ages of the samples are given in thousands of years ago (kya).

We ascertained SNPs at SGDP filter level 1, then used the genotype calls at filter level 0 at the ascertained SNPs. When ascertaining SNPs, we limited ourselves to sites that were polymorphic among the samples excluding MA1 and Neanderthal. We excluded MA1 during ascertainment due to its low coverage. We also excluded the Neanderthal sample during ascertainment because it had substantially fewer new mutations than expected based on its age; it was unclear whether this was due to changes in the mutation rate on that lineage since its deep split with modern humans, or whether this was an artifact of the SNP calling strategy used by SGDP. We used Corollary 1 to correct the normalizing constant of the SFS due to excluding MA1 and Neanderthal during ascertainment; in particular, we normalized the SFS by the total branch length of the subtree excluding MA1 and Neanderthal.

To avoid biases in ancient DNA caused by deamination (Dabney et al., 2013), we removed all transitions (i.e. A↔G and C↔T mutations), keeping only the transversions. We used Chimp as a proxy for the ancestral allele, removing all sites where the Chimp allele was missing. After data cleaning, we were left with 2,444,888 autosomal transversion SNPs that were segregating among the samples excluding MA1 and Neanderthal.

#### A.3.2 Model fitting procedure

We fit the model in Figure 2 in an iterative fashion, adding populations in one at a time and re-estimating the parameters. We started with a tree including the 4 populations Mbuti, Loschbour, Han, and Ust’Ishim, with no admixture events. This initial model had 8 parameters: the population sizes at the Han, Mbuti, and Loschbour leaves, the population size at the Eurasian and human ancestor (the Ust’Ishim leaf was set to the ancestral Eurasian population size), and the times that the Han, Ust’Ishim, and Mbuti populations diverged from Loschbour.

We next added in the Neanderthal population, with an admixture event from Neanderthal to the Eurasian ancestor. We added parameters for the Neanderthal-human split time, the Neanderthal-Eurasian introgression tie and strength, the ancestral Neanderthal-human population size, and a Neanderthal population exponential decline rate starting at the Eurasian-Mbuti split, following results from Prüfer et al. (2014).

We followed by adding on the MA1 sample, adding a single parameter for its split time (we fixed its population size to be the ancestral Eurasian population size). We then added the LBK early farmer, adding 4 parameters for its divergence time, Basal Eurasian admixture time and strength, and the split time of the Basal Eurasian lineage. Finally, we added in a Sardinian population, along with parameters for its population size, its split time, and admixture times and strength from the Loschbour WHG.

At each step, we re-estimated all parameters using the L-BFGS-B optimization algorithm, maximizing the multinomial composite likelihood (11). For the final few models (adding in LBK and then Sardinian) we initialized each estimation with a stochastic gradient descent before finishing with L-BFGS-B.

#### A.3.3 Mutation rate estimation

We used the within-population nucleotide diversity to estimate the mutation rate. The nucleotide diversity is the number of sites where 2 random alleles (drawn without replacement) differ. The empirical value of the nucleotide diversity in population *i* is

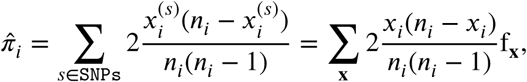

while the expected value of the nucleotide diversity per unit mutation rate is

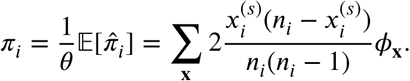

This yields a mutation rate estimate *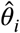* for each population, given by *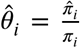*. We then averaged over *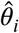* obtain our combined estimate *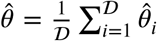*. To obtain the per-site mutation rate we need to divide by the number of bases *L* after our data cleaning process. We approximated this as follows: at SGDP filter level 1, there are about 2.13 × 10^9^ sites per individual; multiply this by 0.93 due to excluding the sex chromosomes; multiply this by 0.32 due to only using transversions; finally, multiply by 0.93 to account for excluding sites missing the Chimp allele, and multiply by 1.1 to account for using genotype calls at filter level 0. The latter 2 factors (accounting for sites missing the Chimp allele and for adding in genotype calls at filter level 0) we estimated by observing how the number of heterozygotes per individual changed after these data cleaning steps.

## Acknowledgments

This research is supported in part by an NIH grant R01-GM109454, a Packard Fellowship for Science and Engineering, the Human Frontiers Science Program LT000402/2017, and Wellcome Trust grants WT206194 and RG89781. YSS is a Chan Zuckerberg Biohub investigator.

## Footnotes

1 We omit certain graph preprocessing steps (moralization and triangulation) from the general junction tree algorithm which are necessary in our setting.

2 The calls to DP(**e**_0_, …, **e**_0_) and 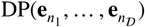 only need to be computed once for all statistics.

